# XGSEA: CROSS-species Gene Set Enrichment Analysis via domain adaptation

**DOI:** 10.1101/2020.07.21.213645

**Authors:** Menglan Cai, Canh Hao Nguyen, Hiroshi Mamitsuka, Limin Li

## Abstract

Gene set enrichment analysis (GSEA) has been widely used to identify gene sets with statistically significant difference between cases and controls against a large gene set. GSEA needs both phenotype labels and expression of genes. However, gene expression are assessed more often for model organisms than minor species. More importantly, gene expression could not be measured under specific conditions for human, due to high healthy risk of direct experiments, such as non-approved treatment or gene knockout, and then often substituted by mouse. Thus predicting enrichment significance (on a phenotype) of a given gene set of a species (target, say human), by using gene expression measured under the same phenotype of the other species (source, say mouse) is a vital and challenging problem, which we call CROSS-species Gene Set Enrichment Problem (XGSEP). For XGSEP, we propose XGSEA (Cross-species Gene Set Enrichment Analysis), with three steps of: 1) running GSEA for a source species to obtain enrichment scores and *p*-values of source gene sets; 2) representing the relation between source and target gene sets by domain adaptation; and 3) using regression to predict *p*-values of target gene sets, based on the representation in 2). We extensively validated XGSEA by using four real data sets under various settings, proving that XGSEA significantly outperformed three baseline methods. A case study of identifying important human pathways for T cell dysfunction and reprogramming from mouse ATAC-Seq data further confirmed the reliability of XGSEA. Source code is available through https://github.com/LiminLi-xjtu/XGSEA

**Author summary:** Gene set enrichment analysis (GSEA) is a powerful tool in the gene sets differential analysis given a ranked gene list. GSEA requires complete data, gene expression with phenotype labels. However, gene expression could not be measured under specific conditions for human, due to high risk of direct experiments, such as non-approved treatment or gene knockout, and then often substituted by mouse. Thus no availability of gene expression leads to more challenging problem, CROSS-species Gene Set Enrichment Problem (XGSEP), in which enrichment significance (on a phenotype) of a given gene set of a species (target, say human) is predicted by using gene expression measured under the same phenotype of the other species (source, say mouse). In this work, we propose XGSEA (Cross-species Gene Set Enrichment Analysis) for XGSEP, with three steps of: 1) GSEA; 2) domain adaptation; and 3) regression. The results of four real data sets and a case study indicate that XGSEA significantly outperformed three baseline methods and confirmed the reliability of XGSEA.

## Introduction

Due to recent advancement of modern experimental technologies, currently we have a massive amount of basic biological data. For example, next-generation sequencing technology has made sequencing faster and lower-cost, generating an incredible number of sequences. This situation makes bioinformatics tools more promising in retrieving biological knowledge from data. For example, gene set enrichment analysis (GSEA) [1] has been well used in biology and related areas, which can rank gene set(s) most relevant (precisely, statistically significant) to binary-labeled gene expression measurement. However, GSEA needs gene expression data labeled binary, such as control and case, and is heavily affected by missing data.

Indeed gene expression are now measured by more speedy and precise techniques like RNA-Seq than cDNA microarray, while measuring gene expression is still costly both on money and time. Existing expression data often has strong bias in measured organisms or species. Model organisms, such as *Mus musculus, Caenorhabditis elegans, Arabidopsis thaliana*, etc., are well measured, while data on minor species are relatively insufficient. Additionally, human gene expression data are unable to be measured under some specific conditions, due to high risk of direct experiments on human, such as non-approved treatment or gene knockout. On the other hand, mouse is usually used to study human disease [2, 3] because of lower cost, lower risk and relatively strong homology relationship with human [4]. However, there exists essential differences between mouse and human [5–8]. Effective treatments developed by mouse data often fail in human clinical trials [9, 10]. Thus it would be strongly expected to develop a method to bridge the gap between expression data of different species, such as human and mouse.

We consider a problem of predicting enrichment significance of given gene sets of one species (such as human) without gene expression, by using sufficient gene expression data of another species (such as mouse). The assumption behind this problem is that both expression data are measured under the same phenotype. We call this problem *cross-species gene set enrichment problem* (XGSEP). Fig 1 shows a schematic picture of the problem setting of XGSEP. Assume that we have enough data behind XGSEP for human and mouse (more generally *target* and *source*), except target expression data. A gene set, either from mouse or human, could be represented as a binary annotation vector with dimension being the number of all genes in the expression data, representing whether the corresponding gene is in the gene set. The enrichment significance (such as *p*-values) of a source gene set *S* with an annotation vector ***x***_*s*_ can be computed by traditional GSEA. The goal of XGSEP is to predict the enrichment significance for a target gene set *T* with an annotation vector ***x***_*t*_, which might have a different dimension from ***x***_*s*_ since the number the total genes for target (human) and source (mouse) are different. Note that the sequence homology between target genes and source genes is assumed to be represented by binary matrix ***M***, which should be important information for the prediction.

**Fig 1.**
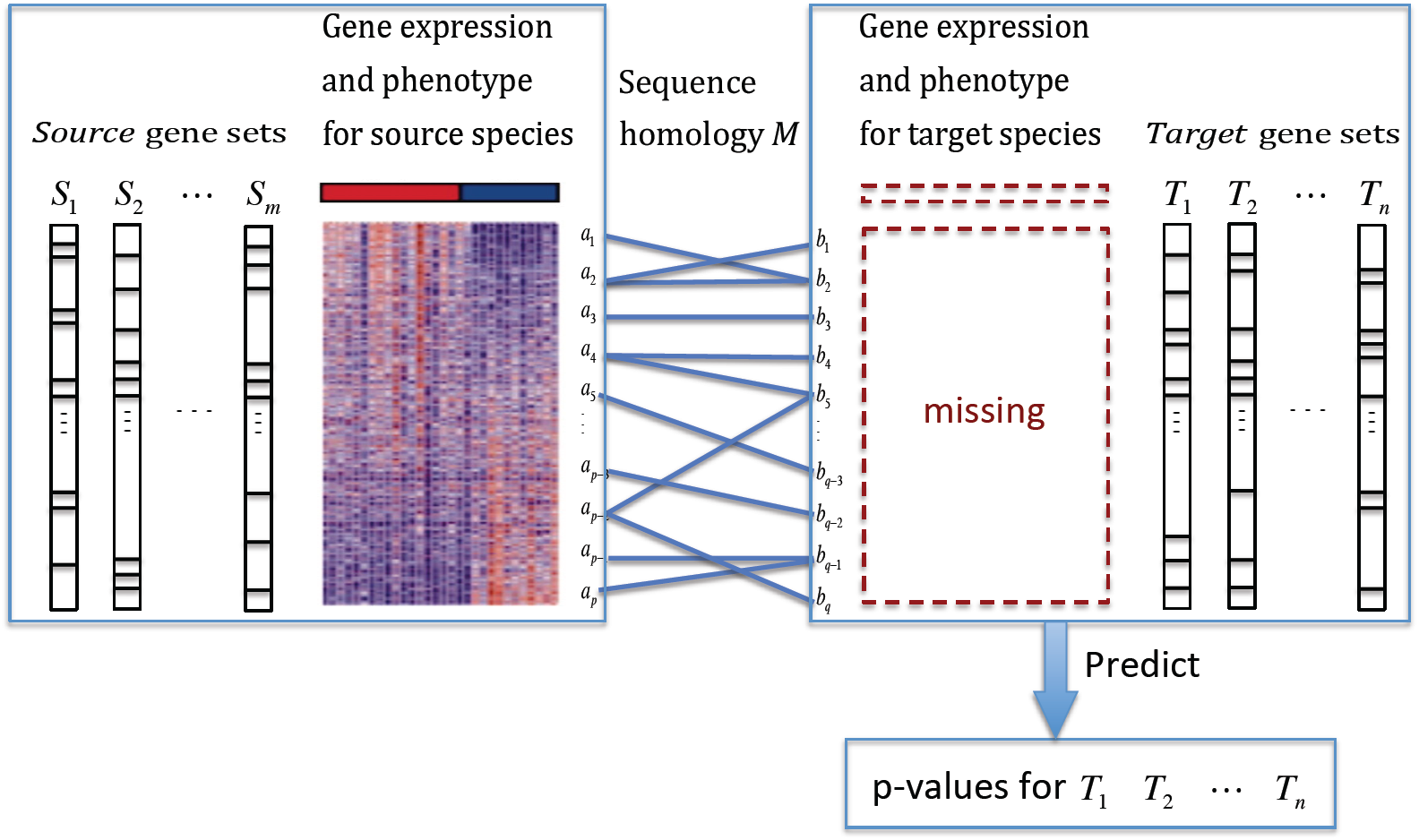
XGSEP: Cross-species gene set enrichment problem, to predict enrichment *p*-values of target gene sets by using source gene sets, gene expression data and sequence homology between target and source genes.

A naive idea for XGSEP would be to first find a source gene set ***x***_*s*_, most homologous to genes in a particular target gene set ***x***_*t*_, by using ***M***. Then GSEA is run over source expression data and ***x***_*s*_. The resultant *p*-value for ***x***_*s*_ is considered as a prediction of the enrichment *p*-value for ***x***_*t*_. The method is simple and fast, but the homology relationship between source and target is often complex, and thus homologous source gene set ***x***_*s*_ cannot be clearly defined. Also using ***M*** directly would be not robust.

Our idea for XGSEP is, rather than focusing on only one gene set, to consider many gene sets at once and train a predictive machine learning model by these gene sets. Suppose that we have source gene sets *S*_1_, *…, S*_*m*_ and target gene sets *T*_1_, *…, T*_*n*_, with annotation matrices 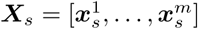 and 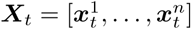, respectively. Then the enrichment *p*-value for the source gene sets can be computed beforehand (by traditional GSEA). The goal of XGSEP is to predict enrichment *p*-values for target gene sets 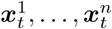. Note that ***X***_*s*_ (training data) and ***X***_*t*_ (test data) are different in size of rows (number of genes), and thus it is difficult to compare the two matrices directly, meaning that a regular machine learning model such as a classifier generated by ***X***_*s*_ cannot be run directly over test data ***X***_*t*_. Thus a further idea is to transform both the target and source species into a common space so that the target and source genes can be compared. However this idea cannot be realized by regular machine learning models by the above problem of difference in size between training and test data. We solve this problem by domain adaptation, transfer learning between two domains: target and source. In general domain adaption, a machine learning model, trained by a larger amount of labeled samples from a source domain, is applied to a target domain with very few or no labeled samples [11]. This is exactly the same situation of XGSEP. A common way of domain adaptation methods is to train a model so that the model can reduce the probability gap between two domains. A possible measure for the probability gap, i.e. the difference of two data distributions, is maximum mean discrepancy (MMD) [12–15]. We will borrow the idea of domain adaptation and MMD to solve XGSEP.

We propose a method, XGSEA, standing for *Cross-species Gene Set Enrichment Analysis*. XGSEA solves XGSEP by three steps: 1) We run GSEA over the source gene sets to obtain gene enrichment scores ***E***_*s*_ and gene enrichment significance ***v***_*s*_. 2) We first define pairwise similarities among gene sets based on ***M***, and then propose a MMD-based domain adaptation method to project ***X***_*s*_ and ***X***_*t*_ into a latent common space with affine mappings ***P***_*s*_ and ***P***_*t*_ to obtain ***Z***_*s*_ and ***Z***_*t*_, respectively, so that i) the probability gap between ***Z***_*s*_ and ***Z***_*t*_ in the latent space is minimized and ii) ***P***_*s*_ and ***P***_*t*_ are smooth over the connection ***M*** between source and target gene sets. By solving this optimization problem, we can obtain the optimal new representations ***Z***_*s*_ and ***Z***_*t*_ for source and target gene sets, respectively. 3) We train a regression model by (***Z***_*s*_, ***E***_*s*_) and run the trained model over ***Z***_*t*_ to predict enrichment scores ***E***_*t*_ for target gene sets and finally *p*-values ***v***_*t*_ with the principle of null hypothesis. Schematically, we may be able to explain our idea by using arrows: 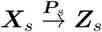 and 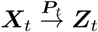, so that the adaptive representations ***Z***_*s*_ and ***Z***_*t*_ for source and target gene sets should have the smallest distribution divergence and preserving their pairwise homology similarities.

The contribution of this work can be summarized into three-fold: 1) We define a problem, XGSEP, which is helpful for understanding a particular phenotype (label) of a species with too limited data to run GSEA. 2) We propose a three-step method called XGSEA for XGSEP through domain adaptation that projects gene sets from two species into a common latent space. This projection is formulated as a nonlinear optimization problem, by which we can estimate the latent space and also estimate the enrichment scores and *p*-values of target gene sets through the latent space. Furthermore the computational complexity of the optimization problem is low enough so that the computation of XGSEA becomes feasible over regular gene annotation matrices. 3) We empirically validated XGSEA by using four different real phenotypes with expression data. The experimental results showed that XGSEA significantly outperformed three baseline methods under various settings. The advantage of XGSEA was further confirmed by a case study of finding significant unknown human pathways for T cell dysfunction and reprogramming from a mouse ATAC-Seq data set.

## Method

To the best of our knowledge, there are no existing work for XGSEP. A similar problem setting might be cross-species gene set analysis (XGSA) [16]. The goal of the XGSA is different with our XGSEP. XGSA aims to compare a gene set from one species with a gene set from another species. That is, XGSA directly examines if two gene sets (from two different species) are significantly different or not, only through the homology between genes in given two gene sets. On the other hand, XGSEP estimates enrichment scores through expression data sets obtained under the same phenotype (see Fig 1, though the target expression is assumed to be missing). Thus XGSA is totally different from XGSEP.

### Problem definition

We have two species, *source* and *target*. Let *A* = {*a*_1_, …, *a*_*p*_} be a source (say mouse) gene set, and *B* = {*b*_1_, …, *b*_*q*_} be a target (say human) gene set. Let ***M***∈ ℝ^*p*×*q*^ be a binary matrix of sequence homology, where the (*i, j*)-element ***M*** (*i, j*) is 1 if source gene *a*_*i*_ is homologous to target gene *b*_*j*_; otherwise zero. Suppose that we have gene expression matrix ***G***_*s*_ with phenotype vector ***y***_*s*_ for source genes only, meaning that we can run GSEA over ***G***_*s*_ and ***y***_*s*_ to compute gene set enrichment significance for an arbitrary source gene set.

Suppose further that we have multiple gene sets for both source and target. Let 𝒮 = {*S*_1_, …, *S*_*m*_} be *m* source gene sets and 𝒯 = {*T*_1_, …, *T*_*n*_} be *n* target gene sets. Thus we have a binary matrix (which we call *annotation matrix*) between *A* (for rows) and 𝒮 (for columns), where 1 means that the corresponding gene is in a gene set; otherwise zero. This can be also for the target side. Let 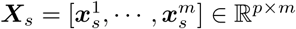 be the annotation matrix for source gene sets *S*_1_, …, *S*_*m*_, where the *i*-th element of 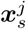 is 1 if gene *a*_*i*_ is in gene set *S*_*j*_; otherwise zero. Similarly, let 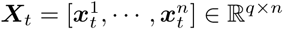 be the annotation matrix for target, where the *i*-th element of 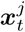 is 1 if gene *b*_*i*_ is in gene set *T*_*j*_; otherwise zero. Then the problem, XGSEP standing for CROSS-species Geneset Enrichment Problem, is, given ***G***_*s*_, ***y***_*s*_, ***X***_*s*_, ***X***_*t*_ and ***M***, to estimate the enrichment *p*-value of each gene set in 𝒯 with respect to the same phenotype of ***y***_*s*_. We propose our method XGSEA, standing for CROSS-species Gene Set Enrichment Analysis, to solve XGSEP by using three steps. Fig 2 shows a schematic picture of the three-step procedure of XGSEA. Below we will explain each of these three steps in detail.

**Fig 2.**
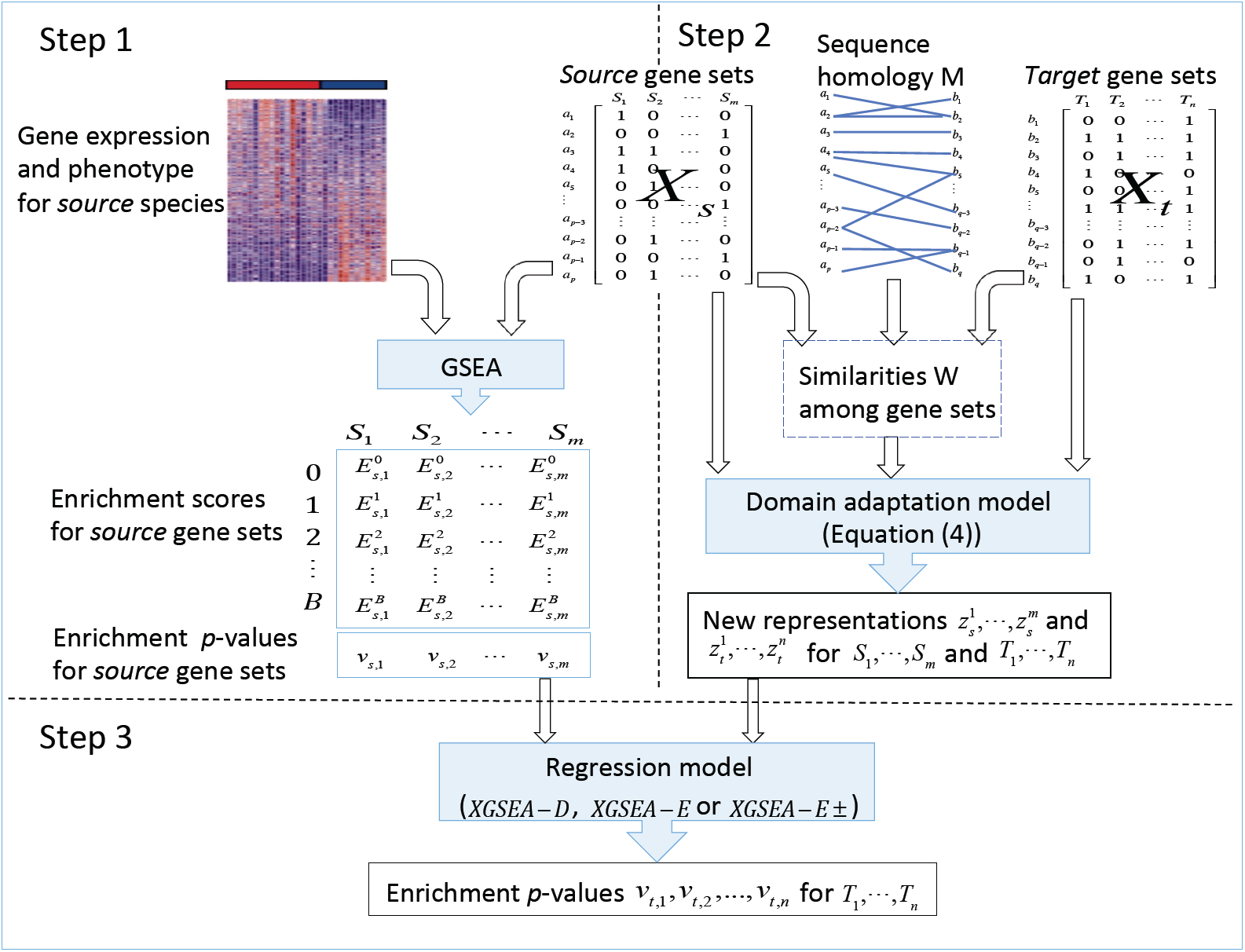
Flow chart of XGSEA: we 1) compute *B*+1 enrichment scores and *p*-values for each source gene set by GSEA, where *B* is the number of permutation, 2) obtain new representations for all source and target gene sets by domain adaptation, and 3) predict enrichment *p*-values for target gene sets by a regression model based on the new representations.

### Step 1: Gene set enrichment analysis for source

Since gene expression ***G***_*s*_ and phenotype ***y***_*s*_ are both available for the source side, we can directly use regular GSEA to obtain *p*-values, *v*_*s*,1_, …, *v*_*s,m*_ for *S*_1_, …, *S*_*m*_, respectively. In fact, *p*-value *v*_*s,i*_ corresponds to null hypothesis 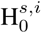: gene set *S*_*i*_ has no association with phenotype ***y***_*s*_ (against the entire set of genes) and can be computed by the following procedure [1].

**1a.** Compute enrichment score 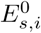 for gene set *S*_*i*_ by using gene expression ***G***_*s*_ and phenotype ***y***_*s*_.

**1b.** Permute the entries in ***y***_*s*_ and recompute the enrichment score for gene set *S*_*i*_. Repeat this step *B* times to generate an empirical null distribution of the enrichment score: E_NULL_ with 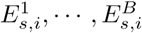.

**1c.** Compute empirical, nominal *p*-value *v*_*s,i*_ for *S*_*i*_ from null distribution E_NULL_ by using the positive (or negative) region of the distribution corresponding to observed enrichment score 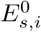.

For source gene set *S*_*i*_, we can compute *B*+1 enrichment scores 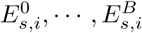 in **1a** and **1b** to compute *p*-value *v*_*s,i*_ in **1c**. Similarly for target gene set *T*_*j*_, we can first predict *B*+1 enrichment scores 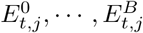 for target gene set *T*_*j*_ and then *p*-value *v*_*t,j*_ in **1c**.

### Step 2: Domain adaptation for source and target gene sets

We project the target and source genes into a common space, to maximally use the information from the source gene side for the target gene sets.

#### Formulating the objective function

We project ***X***_*s*_ and ***X***_*t*_ to a common subspace in ℝ^*d*^ by using affine mappings ***P***_*s*_ ∈ ℝ^*p*×*d*^ and ***P***_*t*_ ∈ ℝ^*q*×*d*^, respectively, such that 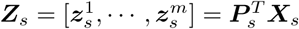 and 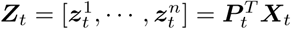.

In this process, we can set the following two reasonable objectives:

1. Probability divergence between ***Z***_*s*_ and ***Z***_*t*_ should be small.
2. Pairwise distances among the gene sets in ***Z***_*s*_ and ***Z***_*t*_ should be preserved.

For the first objective, we use maximum mean discrepancy (MMD) [12, 14]. to measure the divergence. An empirical estimate of MMD can be defined as follows:

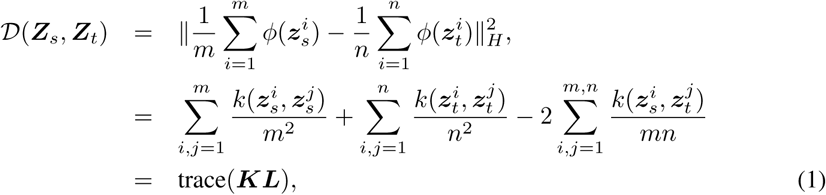

where *ϕ*(·) is a mapping to reproducible kernel Hilbert space *H, k*(·,·) = (*ϕ*(·), *ϕ*(·)) is the kernel associated to this mapping, and

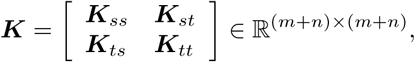

where the (*i, j*)-element of ***K***_*ab*_ is

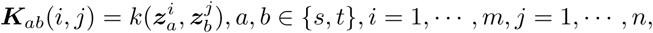

and the (*i, j*)-element of ***L*** is

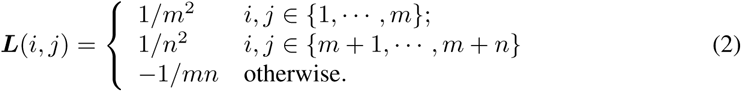

For the second objective, we can first define the pairwise homologous similarity between source gene sets *S*_1_, …, *S*_*m*_ and target gene sets *T*_1_, …, *T*_*n*_ from given data directly as follows,

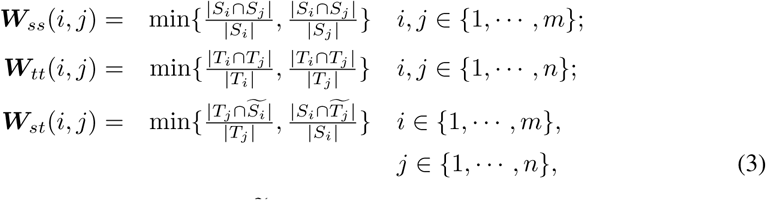

where |*A*| is the number of genes in set 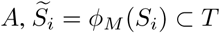 is the set with the target genes homologous to the source genes in *S*_*i*_, and 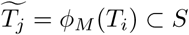 is the set with the source genes homologous to the target genes in *T*_*j*_. The projection should be smooth over homologous similarity matrix 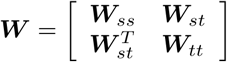.

Thus entirely divergence 𝒟 in (1) should be minimized, being regularized by the smoothness (of the projection) over similarity matrix ***W***. Overall the objective function can be given as follows:

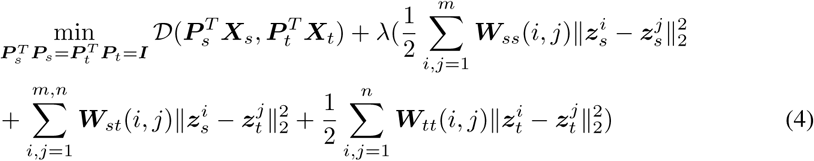

#### Optimization on Grassman manifold

We can use

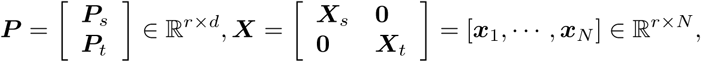

where *r* = *p* + *q*, and *N* = *m* + *n* to write ***Z*** = [***Z***_*s*_ ***Z***_*t*_] = ***P*** ^*T*^ ***X*** ∈ ℝ^*d*×*N*^. Then the first term in (4) can be written as

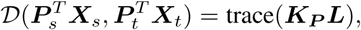

where 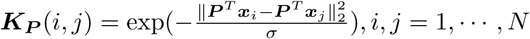, *i, j* = 1, …, *N*, and ***L*** is defined in (2). Note that ***K***_***P***_ depends on ***P***.

Also the regularization term in (4) can be written as

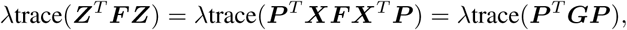

where ***F*** = ***D*** − ***W*** is a Laplacian matrix, ***D*** is a diagonal matrix with ***D***_*ii*_ =∑ _*j*_ ***W***_*ij*_, and ***G*** = ***XF X***^*T*^.

The constraints can be changed from 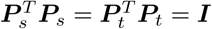 to 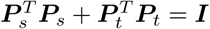 which can avoid that all samples collapse to the origin. Finally (4) can be transformed into an easily understandable form:

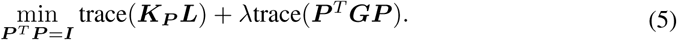

Let *f* (***P***) = trace(***K***_*P*_ ***L***) + *λ*trace(***P*** ^*T*^ ***GP***). The optimization problem 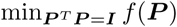 can be solved on the Grassmann manifold, with all linear *d*-dimensional subspaces in ℝ^*p*^, since optimizing *f* (***P***) is not affected by any orthogonal transformation of ***P***. We use the conjugate gradient (CG) algorithm on the Grassmann manifold [17] to solve the optimization problem 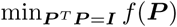. The key step is to compute partial derivative *∂*_***P***_ *f* (***P***), which is used for computing gradient ∇_***P***_ *f* (***P***) of *f* on the manifold at the current estimate ***P*** by ∇_***P***_ *f* (***P***) = *∂*_***P***_ *f* (***P***) − ***PP*** ^*T*^ *∂*_***P***_ *f* (***P***). The search direction is determined at each step by combining the previous search direction with ∇_***P***_ *f* (***P***), and in this direction, a line search along the geodesic at the current estimated ***P*** is performed. Note that partial derivative *∂*_***P***_ *f* (***P***) at the current ***P*** can be obtained as follows

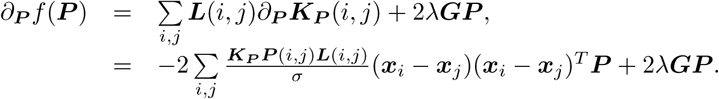

Table 1 shows a pseudocode of the optimization algorithm of Step 2.

**Table 1.**
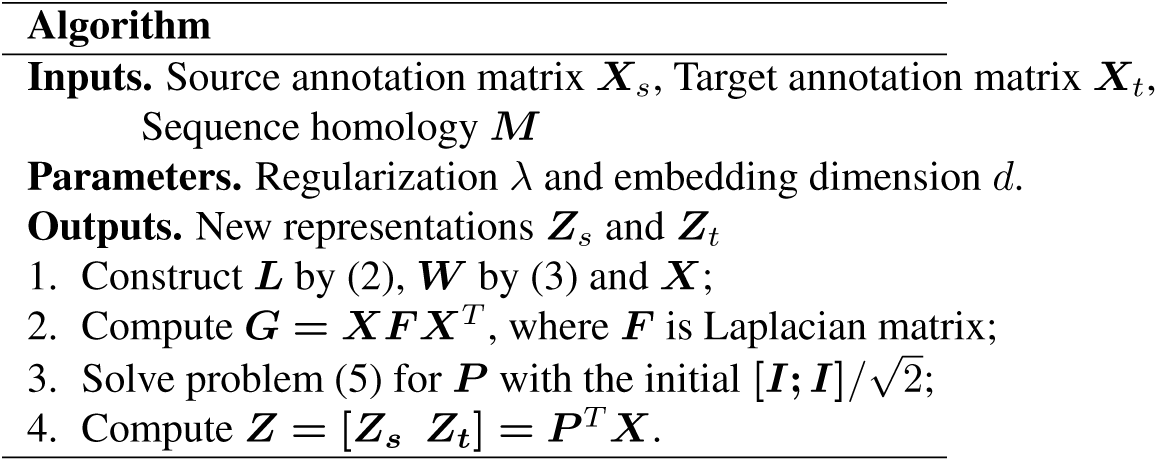
Pseudocode of the optimization algorithm in Step 2 of XGSEA

### Reducing computational complexity

The computational complexity of the above algorithm for solving the optimization problem (5) is *O*(*N* ^2^ + *r*^2^ + *Nrd*). The total number of either human or mouse genes is large, leading to *r*(= *p* + *q*) ≫ *N* (= *m* + *n*). This (large *r*) problem can be a bottleneck for our algorithm, and thus we need to reduce the *r*-related part of this complexity. For this purpose we propose an approach, which uses QR decomposition, by which the computational complexity is reduced to *O*(*N* ^2^). Below, we will explain more detailed manner of our approach.

We first use QR decomposition: ***X*** = ***QR***, where ***Q*** ∈ ℝ^*r*×*N*^ is an orthonormal matrix and ***R***∈ ℝ^*N*×*N*^ is an upper diagonal matrix. By introducing 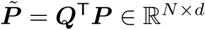, the objective function in (5) can be transformed as follows:

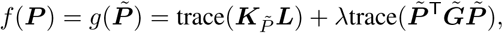

where 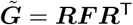, and 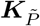 can be obtained using ***R*** since 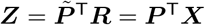. Thus we can first solve a small-scale optimization problem of 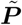, i.e.

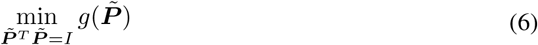

and then obtain the projections 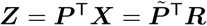. Note that now solving (6) by the above algorithm on the Grassmann manifold needs the computational complexity of only *O*(*N* ^2^).

### Step 3: Enrichment scores and *p*-values for target

The final step in XGSEA is to estimate the *p*-values for the target gene sets, based on the adaptive representations **Z**_*s*_ and **Z**_*t*_ for the source and target gene sets obtained in the above step. One idea is to regress *p*-values on the new representations of the gene sets by logistic regression (XGSEA-D). However, the resulting *p*-values may not obey the principle of null hypothesis. By the principle of null hypothesis, *p*-values is defined as the probability of obtaining the same or more extreme statistics than the observation under null hypothesis, and should be determined by the observed enrichment scores and the null distribution of enrichment scores. Thus another idea is to first predict the observed and null enrichment scores, and then determine *p*-values by its definition. This means that we have one more step to reach *p*-values from the enrichment scores ***E***_*t*_. In detail, the second idea to predict *p*-value for target gene set *T*_*j*_ is that we first predict the enrichment scores *B* + 1 enrichment scores 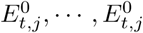 for target gene set *T*_*j*_ and then estimate *p*-value *v*_*t,j*_ by step **1c** in the section of Step1. Based on this idea, we propose to use two regression methods (XGSEA-E and XGSEA-E±). We explain the three methods as follow.

- XGSEA-D: Logistic regression on *p*-values We first train the regression parameters *α* by source gene sets in the following logistic regression model

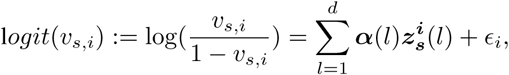

for *i* = 1, …, *m*, and then predict *p*-values for the target gene sets by

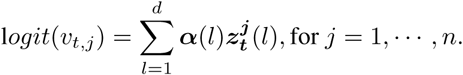

Finally we can obtain the *p*-values of the target species by the following,

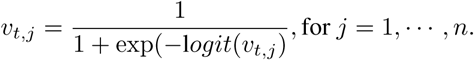
- XGSEA-E: linear regression on enrichment scores Note that we have computed 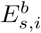, the enrichment score of source gene set *S*_*i*_ at the *b*-th permutation (*b* = 0 means no permutation), in step 1. For any *b* ∈ {0, …, *B*}, we regress the enrichment scores on the new representations of gene sets. The parameter in the regression model can be learnt based on the source gene sets as follows

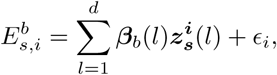

for *i* = 1, …, *m*, and then predict the enrichment scores for target gene sets by

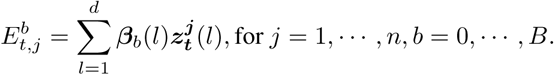

Finally, we can compute the enrichment *p*-values for target gene sets by using step **1c** in the section of Step1.
- XGSEA-E± : linear regression on positive and negative enrichment scores, respectively Similar to XGSEA-E, we predict *p*-values by first estimating enrichment scores for target source gene sets. Different with XGSEA-E, in XGSEA-E± we learn two linear regression models for positive and negative enrichment scores, separately as follows

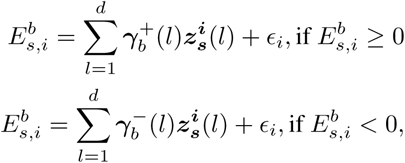

for *i* = 1, …, *m*. The parameters ***γ***^+^ and ***γ***^−^ are learnt by training the source gene sets, and then used to predict enrichment scores for target gene sets by

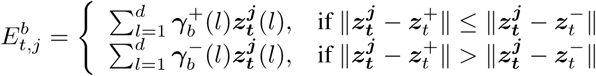

where 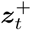 and 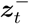 are the centers for ***Z***_*s*_ with positive and negative enrichment scores respectively, and *j* = 1, …, *n, b* = 0, …, *B*.

## Results

### Comparison methods

Since there are no existing methods for XGSEP, we compared XGSEA with three simpler methods, HM_1_, HM_*A*_, and HM_*O*_, which all directly map each target gene to source genes based on sequence homology, and estimate the enrichment *p*-value of target gene set *T* from enrichment *p*-values of particular source gene set *S*. However these three baseline methods take different strategies to generate *S*:

HM_1_: *S* has a randomly chosen gene homologous to each gene in *T* (i.e. |*S*| = |*T* |).

HM_*A*_: *S* has all genes homologous to each gene in *T* (i.e. |*S*| ≥ |*T* |).

HM_*O*_: *S* has, out of gene sets predefined by biological pathways and GO terms, the set with genes most overlapped with those in *T*.

Since we propose three methods, thus we compared totally six methods : XGSEA-D, XGSEA-E, XGSEA-E±, HM_1_, HM_*A*_ and HM_*O*_.

### Data sets

To evaluate the performance of XGSEA, we need target expression data, so that we can compute ground-truth enrichment *p*-values. We collected four gene expression data sets as below, where each data set consists of human (target) and another species (source: mouse or zebrafish) which share the same phenotype. Table 2 shows the statistics of the four data sets.

**Table 2.** Statistics of expression data and human gene sets (𝒯 ^(3)^, where the cutoff for *p*-values was 0.01 and 0.05 for embryonic development and the others, respectively).

| Data set | Species | Gene expression |  |  | Test sets in human |  |
| --- | --- | --- | --- | --- | --- | --- |
|  |  | #genes | #samples | #labels | #sets | #positive sets |
| Embryonic | Human | 14,766 | 29 | 7 | 674 | 24 |
|  | Mouse | 13,879 | 17 | 6 |  |  |
| Brain | Human | 44,030 | 12 | 2 | 674 | 24 |
|  | Mouse | 9,653 | 8 | 2 |  |  |
| Ovarian | Human | 21,188 | 17 | 2 | 674 | 13 |
|  | Mouse | 45,101 | 11 | 2 |  |  |
| Melanomas | Human | 42,346 | 12 | 2 | 664 | 15 |
|  | Zebrafish | 13,620 | 8 | 2 |  |  |

- **Embryonic Development** (human and mouse): The two datasets were collected from www.ncbi.nlm.nih.gov/geo with accessing number GSE44183. Both gene expression datasets were obtained from single cell RNA sequencing. In the human dataset, there are 29 samples with 14,766 genes and seven embryonic development stages, oocytes, pronucleus, zygote, 2-cell, 4-cell, 8-cell and morula. For the mouse, there are 17 samples with gene expression levels of 13,879 genes at sixembryonic development stages, oocytes, pronucleus,2-cell, 4-cell, 8-cell and morula. These datasets were used in a cross-species study [18] already, while this study is not on GSEA.
- **Brain Cancer** (human and mouse): The datasets of the two species were downloaded from GEO with accession number GSE45874 and GSE38591, respectively. Both datasets were measured by microarray. The human dataset has 44,030 genes with six disease and six control samples, while the mouse dataset has 9,653 genes with four disease and four control samples. These datasets were also used in another cross-species study [19], while this study is also not on GSEA at all.
- **Ovarian Cancer** (human and mouse): The two Microarray gene expression datasets were downloaded from GEO with accession number GSE6008 and GSE5987, respectively. The human dataset has 21,188 genes with thirteen mucinous ovarian tumors and four control samples, while the mouse dataset has 45,101 genes with seven disease and four control samples. These datasets were also used in the cross-species study [19].
- **Melanomas** (human and zebrafish): The Microarray gene expression datasets of the two species were downloaded from GEO with accession number GSE83343 and GSE83399, respectively. The human dataset has 42, 346 genes with eight disease and four control samples, while the zebrafish dataset has 13, 620 genes with five disease and three control samples. These datasets were collected from two different studies [20, 21].

We then accessed Ensembl BioMart through http://www.ensembl.org/ [22] to retrieve homology relationships between 19,404 human and 19,614 mouse genes, and also 16,070 human and 18,324 zebrafish genes. The homology data from Ensembl is produced at the protein level rather than the DNA level by whole-genome alignments of vertebrate species [23, 24]. Fig 3 shows two homology matrices between human and mouse (left) and between human and zebrafish (right).

**Fig 3.**
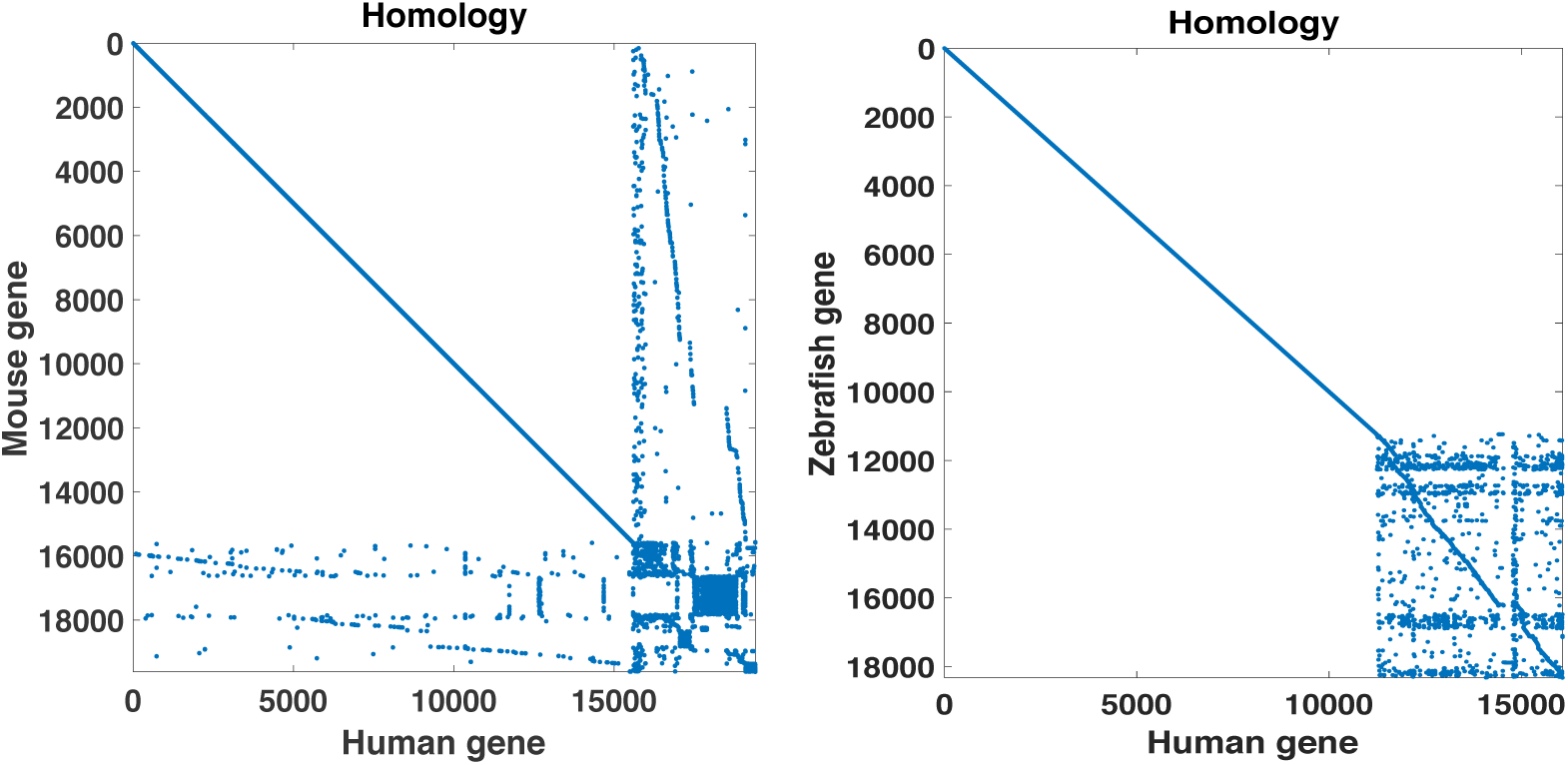
Homology relationships for (left) mouse-human and (right) zebrafish-human.

We can see that genes cannot be assigned in a simple one-to-one correspondences manner. We collected 674 human gene sets (pathways) from Reactome in Molecular Signatures Database (MSigDB), 2,250 mouse gene sets from http://baderlab.org/GeneSets and 1,550 zebrafish gene sets from http://bioinformatics.org/go2msig/.

### Experimental setting

In our experiments, we take human species as target species, and take mouse or zebrafish as the target species. We apply our XGSEA approach to predict the enrichment p-values for the 674 human pathway gene sets 𝒯 = {*T*_1_, …, *T*_*n*_} (*n* = 674), for embryonic development and brain, ovarian and melanomas, respectively. For the target gene sets 𝒯 = {*T*_1_, …, *T*_*n*_}, we take the training source gene sets 𝒮 = {*S*_1_, …, *S*_*n*_} in the XGSEA, where *S*_*i*_ corresponds to *T*_*i*_, meaning that each gene in *S*_*i*_ is homologous to one or more genes in *T*_*i*_.

To sufficiently evaluate our XGESA method, we predict enrichment p-values for target gene sets with three experimental settings. Note that the homology between two genes can be classified into four types: one-to-one, many-to-one, one-to-many, and many-to-many, where one-to-one means only one gene in one side is homologous to only one gene in the other side. First level is for simple target gene sets 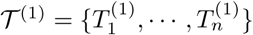, where each 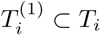 only includes the target genes in *T*_*i*_ with label ‘one-to-one’. For this case, each target gene *g* in set 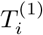 only have one homologous source gene, which does not have any other homologous target gene except *g*. The second case is for more complex target gene sets 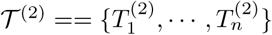, where each 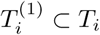 only includes the target genes in *T*_*i*_ with label ‘one-to-one’ and ‘one-to-many’. For this case, each target gene *g* in set 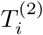 only have one homologous source gene, which may or may not have other homologous target genes besides *g*. The third case is the most complicated case with pathway target gene sets 𝒯 ^3^ = 𝒯 = {*T*_1_, …, *T*_*n*_}, where the target genes may have any of four labels.

In summary, we consider three levels for 𝒯, i.e. 𝒯 ^(1)^, 𝒯 ^(2)^ and 𝒯 ^(3)^, where 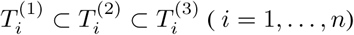

𝒯 ^(1)^ (simple): each set in 𝒯 ^(1)^ has one-to-one genes only. That is, target gene 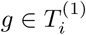 has only one homologous source gene *s*, which has no other homologous target genes except *g*.

𝒯 ^(2)^ (medium): each set in 𝒯 ^(2)^ has one-to-one or many-to-one genes. That is, target gene 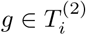 has always only one homologous source gene *s*, which has one or more homologous target genes including *g*.

𝒯 ^(3)^ (complex): each set in 𝒯 ^(3)^ target gene *g* may have one or more homologous source gene, and one of them *s* also may have one or more homologous target gene, including *g*.

### Evaluating XGSEA by supervised learning

Each target gene set has a ground-truth *p*-value. In evaluation, target gene sets with smaller true *p*-values should be predicted to have smaller *p*-values. In this light, we examined XGSEA and competing methods in a supervised manner: we set a cutoff (significance level) for the ground-truth *p*-values of target gene sets so that a gene set is a positive instance if the true *p*-value of this instance is lower than the cutoff; otherwise a negative. This means that we can control the number of positives (and negatives) by changing the cutoff. Then once after true positives (and true negatives) are determined by the cutoff for *p*-values in the above manner, we can examine the ROC (receiver operator characteristics) curve (and also precision-recall (PR) curve) by sorting the predicted *p*-values for gene sets in the ascending order. Note that this is regular validation of supervised learning (more precisely binary classification).

The *d* and *λ* were chosen from {5, 10, 20, 30, 40, 50} and {0.01, 0.1, 1, 10, 100}, respectively, to give the best performance under each experimental setting.

### Performance on four real data sets

Fig 4 shows sample ROC and PR curves for one of the four data sets, i.e. embryonic development under 𝒯 ^(3)^ with the cutoff (for *p*-values) of 0.01. These figures shows that XGSEA (red and blue) look outperformed compared naive methods (green, yellow and light blue), except for XGSEA-D (black), indicating that regression of *p*-values on *p*-values directly may perform badly, as we expected. Note that there exist overlaps between XGSEA and naive methods, making the comparison unclear. Thus we checked the performance difference more systematically.

**Fig 4.**
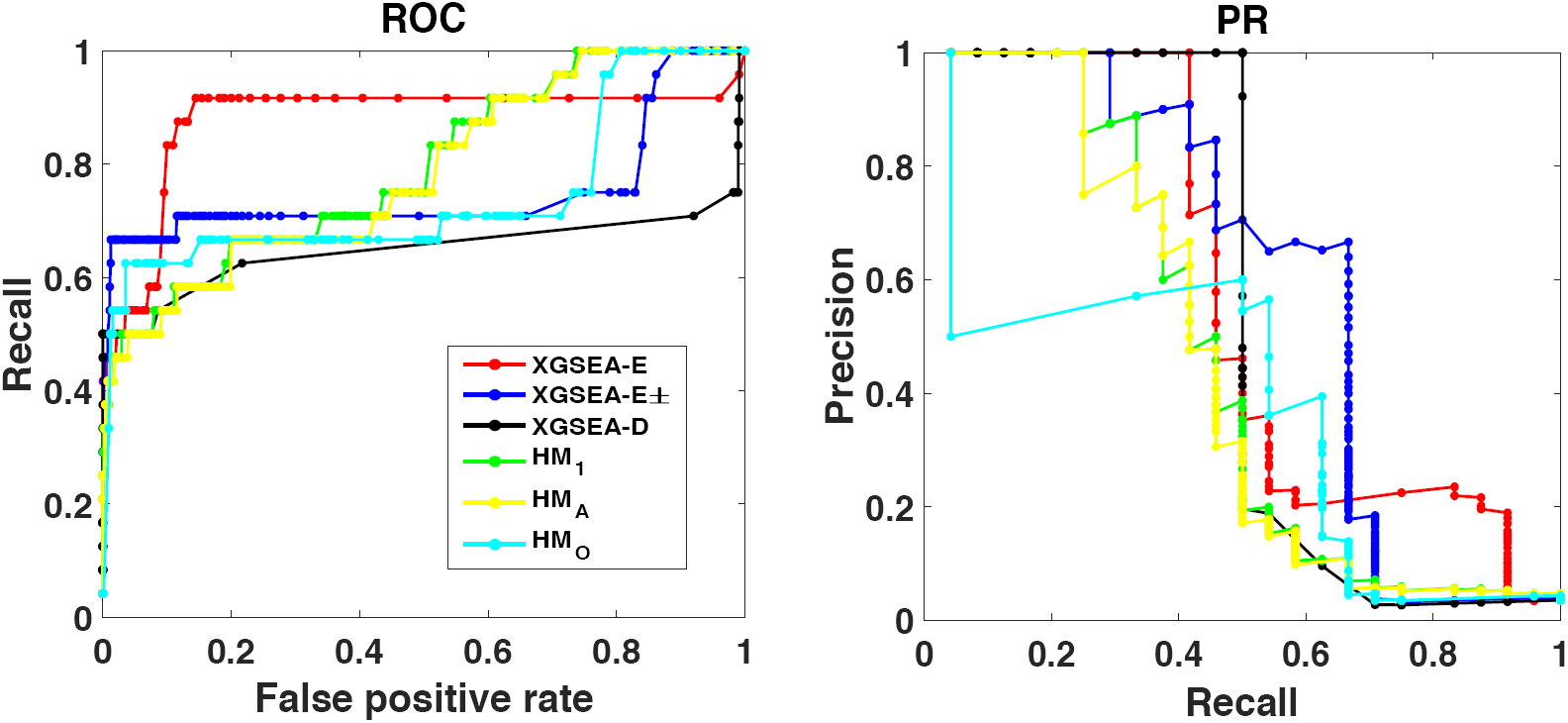
ROC and PR curves by XGSEA-D (black), XGSEA-E (red), XGSEA-E± (blue), HM_1_ (green), HM_*A*_ (yellow) and HM_*O*_ (light blue) for embryonic deveopment under 𝒯 ^(3)^.

We changed the cutoff for *p*-values: {5*e*-1, 1*e*-1, 5*e*-2, 2.5*e*-2, 1*e*-2, 5*e*-3, 2.5*e*-3, 1*e*-3}, resulting in changing the number of true (ground-truth) positives. That is, the number of true positives becomes smaller for smaller cutoff values. Fig 5 (left column) shows, changing the cutoff for *p*-values, the AUC (area under the ROC curve) of all competing methods on all four real data sets under 𝒯 ^(3)^. The AUC increased as the cutoff was decreasing (the number of true positives was decreasing). For most of the changing cutoff values, XGSEA (black, red and blue) showed better AUCs than the three baseline methods (green, yellow and light blue).

**Fig 5.**
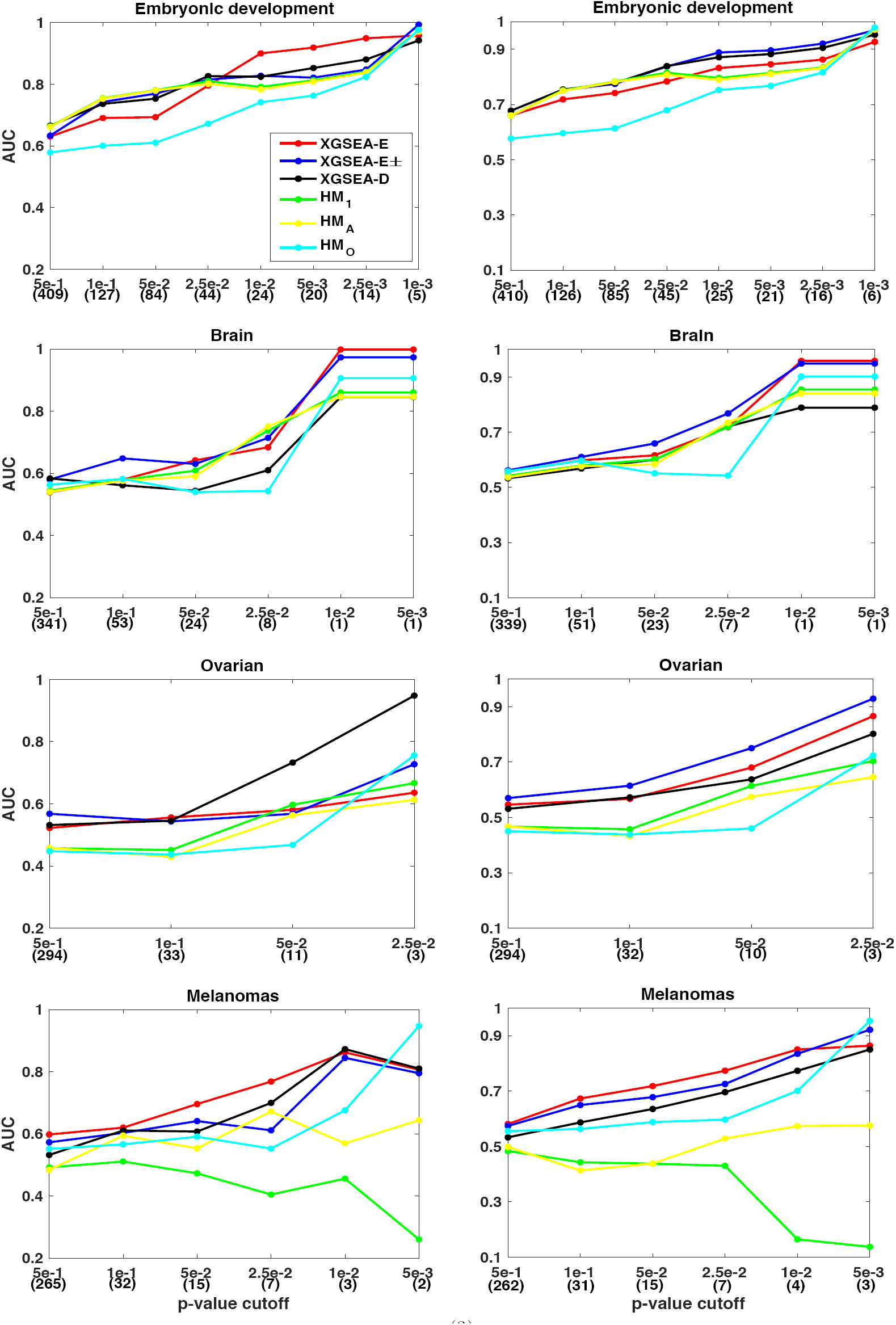
(Left column) AUCs on four data sets (𝒯 ^(3)^), changing the cutoff for *p*-values. (Right column) Bootstrapped (20 trials) AUCs under the same condition as the left column. Compared methods are XGSEA-D (black), XGSEA-E (red), XGSEA-E± (blue), HM_1_ (green), HM_*A*_ (yellow) and HM_*O*_ (light blue).

### Stabilized results by bootstrapping

Smaller cutoff values, such as 5*e*-3, resulted in an extremely few number of positives. For example, brain cancer had only one positive for the cutoff of 5*e*-3. Also each AUC (in the left column of Fig 5) was obtained by only one trial of training and test. These two aspects made AUCs in the left column of Fig 5 rather unstable. To resolve this issue, we conducted bootstrapping on 674 human gene sets of 𝒯 ^(3)^ by repeating sampling with replacement 20 times, resulting in 20 AUCs, which were averaged. Fig 5 (right column) shows the averaged AUCs (over 20 trials) of all methods on all four real data sets, under 𝒯 ^(3)^, changing the cutoffs for *p*-values. Comparing with the left column, the results were stabilized, clarifying the performance advantage of XGSEA (black, red and blue) over the three baseline methods (green, yellow and light blue). In particular, even the difference between the three proposed methods became clearer.

We then, fixing the cutoff value, examined the performance of the competing methods. Table 3 shows (bootstrapped) AUCs under three different gene sets (𝒯 ^(1)^, 𝒯 ^(3)^ and 𝒯 ^(3)^) by all six methods, fixing the cutoff at 0.01 for embryonic development and 0.05 for the other data sets. This table shows that XGSEA significantly outperformed the baseline methods. For example, XGSEA-E± achieved the best in nine out of all 12 cases, followed by XGSEA-E± of three cases. Any naive method could neither be the best nor the second best in all cases, the difference from the best being statistically significant in *t*-test over 20 trials. Also the AUC of 𝒯 ^(1)^ was not necessarily higher than 𝒯 ^(2)^ (also 𝒯 ^(3)^), since each one-to-one homologous gene pair between two species is not necessarily the same gene, which would be prediction-wise harder than the case that the target and source gene sets share the same gene.

**Table 3.** AUCs of six competing methods on four data sets and three target gene sets. The best and second best in each row are in bold and underlined, respectively. The *p*-value by *t*-test between the best and each corresponding naive method is shown in brackets.

| Data set |  | HM<sub>I</sub> | HM<sub>A</sub> | HM<sub>O</sub> | XGSEA-D | XGSEA-E | XGSEA-E± |
| --- | --- | --- | --- | --- | --- | --- | --- |
| Embryonic | $\mathcal{T}^{(1)}$ | 0.81 (6.59e-06) | 0.81 (6.59e-06) | 0.75 (1.48e-08) | <u>0.86</u> | 0.80 | **0.89** |
| | $\mathcal{T}^{(2)}$ | 0.80 (2.12e-06) | 0.80 (2.12e-06) | 0.74 (6.84e-09) | <u>0.86</u> | 0.83 | **0.89** |
| | $\mathcal{T}^{(3)}$ | 0.79 (1.58e-09) | 0.80 (4.43e-09) | 0.75 (3.73e-11) | <u>0.87</u> | 0.83 | **0.90** |
| Brain | $\mathcal{T}^{(1)}$ | 0.66 (3.14e-01) | 0.66 (3.14e-01) | 0.58 (1.14e-05) | 0.60 | **0.68** | <u>0.67</u> |
| | $\mathcal{T}^{(2)}$ | 0.59 (1.00e-04) | 0.59 (1.00e-04) | 0.57 (5.59e-06) | 0.60 | <u>0.66</u> | **0.67** |
| | $\mathcal{T}^{(3)}$ | 0.58 (1.75e-07) | 0.60 (1.36e-05) | 0.55 (2.64e-07) | 0.61 | <u>0.63</u> | **0.68** |
| Ovarian | $\mathcal{T}^{(1)}$ | 0.45 (2.53e-12) | 0.45 (2.53e-12) | 0.57 (1.65e-04) | <u>0.67</u> | 0.64 | **0.70** |
| | $\mathcal{T}^{(2)}$ | 0.56 (6.72e-09) | 0.56 (6.72e-09) | 0.50 (2.07e-08) | 0.67 | <u>0.69</u> | **0.75** |
| | $\mathcal{T}^{(3)}$ | 0.57 (5.60e-12) | 0.61 (1.50e-07) | 0.46 (6.60e-14) | 0.65 | <u>0.70</u> | **0.77** |
| Melanomas | $\mathcal{T}^{(1)}$ | 0.72 (3.65e-12) | 0.72 (3.65e-12) | 0.47 (2.10e-16) | 0.84 | **0.92** | <u>0.87</u> |
| | $\mathcal{T}^{(2)}$ | 0.63 (6.14e-05) | 0.63 (6.14e-05) | 0.48 (8.01e-14) | 0.74 | <u>0.80</u> | **0.81** |
| | $\mathcal{T}^{(3)}$ | 0.44 (1.74e-16) | 0.44 (2.90e-15) | 0.59 (4.68e-06) | 0.64 | **0.72** | <u>0.71</u> |

### Robustness against parameter value change

We examined the performance robustness of XGSEA, regarding parameter (*λ*) variation. Fig 6 shows AUCs of XGSEA-E under three gene sets (𝒯 ^(1)^ (red), 𝒯 ^(2)^ (blue) and 𝒯 ^(3)^ (black)) on embryonic development and melanomas, when *λ* is one of {1*e*-4, 1*e*-3, 1*e*-2, 1*e*-1}. This figure shows that AUC of XGSEA-E was rather stable within the given range, implying that the advantage over the baseline methods will be kept constantly.

**Fig 6.**
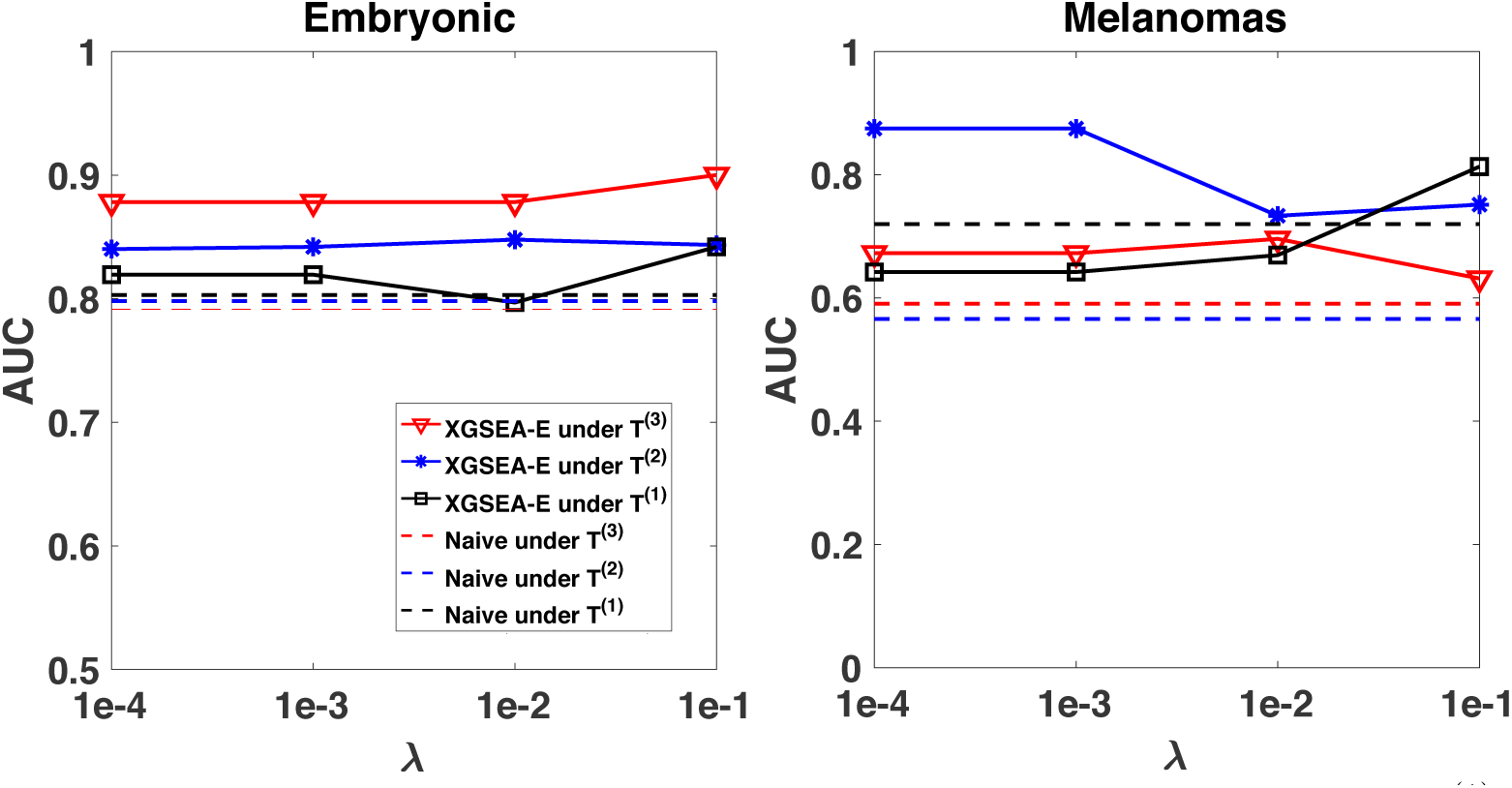
AUCs of XGSEA-E (solid line) and the best of naive methods (dotted line) under 𝒯 ^(1)^ (red), 𝒯 ^(2)^ (blue) and 𝒯 ^(3)^ (black) on (right) embryonic development and (left) melanomas.

### Effect of similarity and homology on predictive performance

We examined the contribution of three types of gene set similarity, i.e. *W*_*ss*_, *W*_*st*_ and *W*_*tt*_, used in XGSEA, by modifying the objective function in the formulation of XGSEA. The objective function of XGSEA is given by (4), which has four terms, where the first term is the divergence and the last three terms are for *W*_*ss*_, *W*_*st*_ and *W*_*tt*_. We then generated four different variants of (4), as follows:

MMD: only divergence, i.e. no terms on gene set similarity.

MMD+W: divergence and two terms on *W*_*ss*_ and *W*_*tt*_.

MMD+B: divergence and the term on *W*_*st*_. MMD+WB: original objective function, i.e. (4).

We applied these four variants to embryonic development data with target gene set 𝒯 ^(3)^. Table 4 shows AUCs obtained with the cutoff (for *p*-values) of 0.01. From Table 4, MMD+WB (i.e. original (4)) achieved the best result for XGSEA-E and XGSEA-E±, and MMD was worst for them. This result implies that all gene set similarity contribute to the performance improvement.

**Table 4.** AUCs of XGSEA in variants and transferabilities, respectively, in embryonic development under gene set 𝒯 ^(3)^.

| | | XGSEA-D | XGSEA-E | XGSEA-E± |
| --- | --- | --- | --- | --- |
| Similarity | MMD | 0.62 | 0.88 | 0.71 |
|  | MMD+W | 0.61 | 0.88 | 0.74 |
|  | MMD+B | 0.62 | 0.80 | 0.73 |
|  | MMD+WB | 0.60 | 0.90 | 0.75 |
| Homology | $M_{50}$ | 0.72 | 0.77 | 0.51 |
| | $M_{500}$ | 0.73 | 0.78 | 0.53 |
| | $M_{5000}$ | 0.75 | 0.84 | 0.73 |
| | $M$ | 0.75 | 0.84 | 0.74 |

We then evaluated the effect of sequence homology on predictive performance, by removing a certain amount of part in sequence homology matrix ***M*** : being motivated by that less homology connectivity between two species would cause poorer performance.

In more detail, we first randomly chose a certain number of genes from the source and target gene sets, respectively, and kept only the part corresponding to these genes in ***M***. Practically we used 50, 500 and 5,000 for this number of selecting genes, resulting in three matrices: ***M***_50_, ***M***_500_ and ***M***_5000_, respectively. Using each of the four sequence homology matrices (including original ***M***), we ran XGSEA over embryonic development data under gene set 𝒯 ^(3)^ to predict enrichment *p*-values.

Table 4 shows the performance results (AUC) of this experiment. The results show that the AUC was reduced by decreasing the number of randomly selected genes, while if the selected number is 5,000, the performance was almost consistent with that of using the original ***M***, implying that interestingly 5,000 genes might be good enough.

### Case study: Identifying human pathways for T cell dysfunction and reprogramming from mouse ATAC-Seq

It is important for cancer immunotherapy to study the epigenetic regulation of T cell dysfunction and therapeutic reprogrammability: a plastic dysfunctional state from which T cells can be rescued, and a fixed dysfunctional state in which cells are resistant to reprogramming [25]. Identifying two (plastic or fixed) dysfunctional chromatin states, through which T cells in tumours differentiate, would be very important to predict, for example, if a patient will respond to a therapy. Using GSE89308 of GEO on ATAC-Seq data of mouse, with 22 samples and the two chromatin states [25], we ran XGSEA-E (*B* = 100,000, *λ*=0.01 and *d*=5) to identify human pathways out of 1,960 Reactome pathways (downloaded from https://reactome.org/download-data).

Table 5 shows 11 human pathways identified by XGSEA-E at the cutoff of 0.05, where the top, “gene expression (transcription)”, and the fourth “immune system” are large pathways with 1367 and 2296 genes, respectively. Obviously due to important chromatin roles in transcription, “gene expression (transcription)” is tightly related to the chromatin states. Also “immune system” definitely plays important roles in T cell dysfunction and reprogramming through a number of membrane proteins, such as CD38, CD101, CD30L, CD5, TCF1, IRF4, BCL2, CD44, PD1, LAG3 and CD62L [25].

**Table 5.** 11 human pathways (with *p*-values) identified by XGSEA-E for T cell dysfunction and reprogramming.

| Pathway | $p$-value |
| --- | --- |
| Gene expression (Transcription) | 0.03 |
| A third proteolytic cleavage releases NICD | 0.03 |
| Signaling by NOTCH | 0.03 |
| Immune System | 0.04 |
| Signaling by NOTCH3 | 0.04 |
| Signaling by NOTCH4 | 0.04 |
| NOTCH2 Activation and Transmission of Signal to the Nucleus | 0.04 |
| Activated NOTCH1 Transmits Signal to the Nucleus | 0.04 |
| Signaling by NOTCH2 | 0.04 |
| Constitutive Signaling by NOTCH1 HD+PEST Domain Mutants | 0.04 |
| Signaling by NOTCH1 | 0.04 |

The remaining 9 pathways are all on Notch signaling pathways, which affect T cells in various ways. Notch signaling pathways play multiple essential roles in thymic T cell development and peripheral T cell differentiation [26]. For example, Delta-like ligand 4 (DLL4) interacts with Notch 1 to specify thymic T cell commitment during lymphocyte development. This Notch pathway regulates CD8+ T cells by directly upregulating mRNA expression of granzyme B and perforin to maintain memory T cells [27]. Furthermore, the Notch pathway plays an important role in antitumor immunity. CD8+ T cell-specific Notch2 deletion impairs antitumor immunity, whereas the stimulation of the Notch pathway can increase tumor suppression. Ezh2, a suppressor of the Notch pathway, regulates effector T cell polyfunctionality and survival by targeting the Notch signaling pathway [28]. Down regulation of Ezh2 could elicit poor antitumor immunity. Besides, Delta-like 1-mediated Notch signaling enhances the conversion of human memory CD4 T cells into FOXP3-expressing regulatory T cells [29]. These facts support the reliability of the pathways identified by XGSEA.

On the other hand, we ran a naive approach, HM_*A*_, over the same data, under the cutoff of 0.05, resulting in 20 pathways showed in Table 6. Although the number of pathways is larger than Table 5, these 20 pathways were diverse and less connected to the chromatin states, such as only two being related to Notch signaling pathways. This result implies that XGSEA-E would be more convincing than HM_*A*_.

**Table 6.** 20 human pathways (with *p*-values) identified by HM_*A*_ for T cell dysfunction and reprogramming.

| Pathway | *p*-value |
| --- | --- |
| Assembly Of The HIV Virion | 4e-5 |
| Membrane binding and targetting of GAG proteins | 2e-4 |
| Mineralocorticoid biosynthesis | 4e-4 |
| Type II Na+/Pi cotransporters | 8e-4 |
| Interactions of Tat with host cellular proteins | 3e-3 |
| Biogenic amines are oxidatively deaminated to aldehydes | 5e-3 |
| SDK interactions | 0.01 |
| Cohesin Loading onto Chromatin | 0.01 |
| XAV939 inhibits tankyrase, stabilizing AXIN | 0.01 |
| Mitotic Telophase/Cytokinesis | 0.02 |
| Interleukin-9 signaling | 0.02 |
| TWIK-related acid-sensitive K+ channel (TASK) | 0.02 |
| Defective Mismatch Repair Associated With MLH1 | 0.02 |
| Budding and maturation of HIV virion | 0.02 |
| CREB phosphorylation through the activation of CaMKK | 0.04 |
| Synthesis of 5-eicosatetraenoic acids | 0.04 |
| Apoptotic execution phase | 0.04 |
| NOTCH2 intracellular domain regulates transcription | 0.05 |
| Toxicity of botulinum toxin type C (BoNT/C) | 0.05 |
| CD209 (DC-SIGN) signaling | 0.05 |

## Conclusion

We have defined XGSEP for promoting GSEA on species with scarce expression data, and proposed XGSEA with three steps, which can be simply: 1) GSEA, 2) domain adaptation, and 3) regression. Our empirical supervised validation over four real data sets revealed that XGSEA outperformed three naive approaches in AUC under various settings, particularly the advantage being proved statistically by bootstrapping and *t*-test. In the case study, mouse ATAC-Seq expression data is used to identify significant human pathways for T cell dysfunction and reprogramming. XGSEA found rather general two pathways related with gene expression (transcription) and immune system, as well as nine Notch signal-related pathways, all being convincing, especially compared with pathways found by a baseline approach.

Improvement of XGSEA would be definitely interesting future work. It would be worth working on exploring a better variation on each of the three steps of XGSEA: Step 1 can be generalized or focused on another statistical problem. Exploring more efficient, robust domain adaptation would be interesting future work for Step 2. Reasonably in Step 3, we can consider more sophisticated regression models. The most key point of XGSEA is Step 2, i.e. domain adaptation, which would be useful for other problems between two species, such as genome wide association studies between a well- and the other less-sequenced species. This direction of applying domain adaptation to various problems would be also promising future work. On the statistical side, we could also further consider the problem of multiple testing and controlling the false discovery rate or family-wise error rate, which have been well studied in regular GSEA.

